# How low can you go? Driving down the DNA input requirements for nanopore sequencing

**DOI:** 10.1101/2021.10.15.464554

**Authors:** Darren Heavens, Darren Chooneea, Michael Giolai, Piotr Cuber, Pia Aanstad, Samuel Martin, Mark Alston, Raju Misra, Matthew D. Clark, Richard M. Leggett

## Abstract

The requirement for large amounts of purified DNA limits many sequencing experiments, especially when seeking to avoid pre-amplification or when using third generation technology to sequence molecules directly. We wanted to test the limits of current nanopore sequencing input requirements and devised a set of experiments to evaluate extraction and library preparation approaches for low inputs.

We found an optimised bead beating approach combined with a magnetic bead protocol, rather than traditional spin columns for DNA extraction, improved both molecule length, integrity score and DNA yield. Through reducing the DNA input to as little as 6.25 % of recommended (25 ng versus 400 ng) and reaction volumes in half, library construction can be completed, and sequencing begun within 20 minutes of sample collection.

Applying these approaches, we demonstrated that our pipeline can be used as a cheap and effective method to *de novo* assemble a genome and identify genes from low quantities and quality of DNA. With our rapid extraction protocol using transportable equipment and low input library construction we were able to generate a de novo assembly from a single insect (*Drosophila melanogaster)* spanning 125 Mbp / 85 % of the reference genome, over 96.9% complete BUSCO genes, with a contig N50 over 1.2 Mbp, including chromosome arm sized contigs, for a modest consumable cost under £600.

## 1. Introduction

Mobile laboratories with the ability to extract DNA, construct libraries and generate sequence data in real time using devices such as Oxford Nanopore Technologies (ONT) MinION are quickly redefining how rapid diagnostics and genome assembly can be deployed in situ. Metabarcoding approaches such as 16S, 18S, ITS, CO1 and MatK analysis are being replaced by whole genome shotgun (WGS) sequencing which offers improved discrimination, particularly for strain level identification^[1,2]^. When optimised, these methods also have the potential to allow gene space assemblies and could become a valuable resource for ambitious efforts such as the Earth BioGenome Project (EBP)^[3]^, where sequencing all eukaryotic species is the aim, and many samples cannot be easily transported to central labs.

There are many challenges which need to be overcome to facilitate robust and universal protocols across a wide range of sample types. For metagenomic applications there can be a mixture of different species present each bringing their own problems for DNA extraction. With Gram-negative bacteria chemical lysis is sufficient to break open cell walls and release DNA but for Gram-positive species spheroplasting with enzymes such as lysozyme is required to puncture the cell wall for effective DNA extraction. Fungal or bacterial spores with tough cell walls typically require bead beating to break them open, however, a commonly perceived disadvantage of bead beating is that molecule length is often compromised as it can be difficult to extract DNA with molecules lengths > 10 Kbp. In some cases this is more than offset by the fact the DNA yields can be considerably greater and are often more representative especially in situations when material is limited such as infant faecal^[4]^ and for low biomass environmental samples like air^[5]^. DNA input requirements can also be out of reach for some non-metagenomic sample types. Fine needle aspirations or core needle biopsies can yield <100ng of material^[6]^ and for these types of samples the ability to sequence native DNA in order to detect base modifications such as methylation can be essential^[7]^, prohibiting the use of amplification strategies such as whole genome amplification (WGA)^[8]^.

ONT protocols are optimised for starting DNA > 30 Kbp and aim to generate read lengths circa 8 Kbp^[9,10]^. These are based on ligation inputs of 1 *μ*g and transposase library requirements of 400 ng. Using these recommended values, we find around 200, 000 reads (or 1-2 Gb) per hour can be generated, though this tails off with time. A major advantage of nanopore sequencing is that DNA molecules of any length can be sequenced. At the recent ONT London Calling meeting it was reported that reads > 4 Mbp had been recorded^[11]^ suggesting that sequencing of a bacterial genome in a single read could be achieved. For metagenomic applications this has led to a focus on DNA extractions to maximise both yield and molecule length and methods such as the three peaks experiment^[12]^ which combines chemical, enzymatic and physical methods to both preserve molecule length and maximise yields. Advances in this field have been driven by the continued identification of new enzymes to break down cell walls. The three peaks experiment utilizes MetaPolyzyme, which is a combination of achromopeptidase, chitinase, lyticase, lysostaphin, lysozyme and mutanolysin, to target many different species^[13]^. A disadvantage of this method is that it takes > 3 hours to complete compared to < 20 minutes for a bead beating and spin column-based protocol such as the Qiagen PowerSoil Pro kit.

Whilst extracting long molecules is important for construction of metagenome assembled genomes (MAGs)^[14-17]^, it isn’t necessarily required for diagnostic applications. With continued improvements in read accuracy, good classification can be achieved with reads around 1-2 Kbp^[4]^ with nanopore technology and there is an advantage to having shorter reads. With pore translocation times of around 450 bp per second, molecules of 1 Kbp in length will have exited a pore in < 3 seconds freeing it up for the next molecule to be sequenced. Therefore, shorter reads are generated faster than longer reads which can help reduce the required sequencing times to make clinical diagnostic decisions.

To further reduce processing time transposases can be used rather than ligation-based techniques for sequencing library construction. Transposase mediated methods can be completed inside 10 minutes, compared to 60 minutes for ligation. For the ONT RAD004 kit chemistry, two one minute incubations, the first at 30 °C to insert the transpose loaded sequence randomly into the DNA and the second, an 80 °C step to denature the enzyme, are followed by a 5 minute incubation at room temperature with a click chemistry enabled adapter. An additional advantage of transposases is that they have the potential to control final molecule length by optimising the ratio of enzyme to DNA which can be useful to target molecules of the optimal size for a given application^[18]^. In addition to this we have previously shown that transposases can be more robust in the presence of a number of different contaminants such as secondary metabolites carried over during DNA extraction^[19]^ These are known to inhibit some enzymatic reactions and ligation methods use more enzymes and enzymatic steps (typically DNA pol I as an end-repair polymerase, a phospho-nucleotide kinase to phosphorylate 5’ nucleotides, either Klenow fragment or a non-proofreading *Taq* polymerase add an A overhang to the 3’ ends of the molecule and a ligase to add an adapter molecule) which could be inhibited.

For genome assembly, it is typically important to sequence an individual of a species rather than pools, to avoid multiple haplotypes complicating assembly graphs. Early NGS projects sequenced bacteria, fresh clonal cultures were grown from single colonies streaked out on a plate^[20]^. The sequencing of an individual mosquito using PacBio data highlighted the assembly benefit compared to previous assemblies obtained from sequencing pools of individuals. Using only 100 ng of DNA for PacBio library construction they achieved a read N50 > 13 Kbp and produced an assembly with a contig N50 > 3.4 Mbp and only 206 contigs^[21]^.

A low input method has also been published that utilises a multi-platform hybrid approach combining Illumina, Nanopore and Hi-C sequencing to assemble a single insect genome with a high degree of contiguity. Working on the *Drosophila* genome, Adams et al. extracted 104 ng of DNA from an individual fly, used a low input transposase method to generate an Illumina library for which they generated 333x coverage, produced an amplified Nanopore library generating 33x coverage with a median read length of 3.5 Kbp and produced 151x coverage of HI-C data. Their final assembly achieved a scaffold N50 of 26 Mbp^[22]^.

To determine assembly accuracy a useful quality control tool to measure gene space completeness is BUSCO analysis which compares the percentage of single copy genes present within an assembly against those expected to be present^[23,24]^. Kingan *et al*. identified 98% of genes as complete and Adams *et al*. 95%.

Simply put it is the combination of the amount of DNA that can be extracted, the length of the molecules sequenced, the amount of sequence generated and the frequency and nature of any repeat elements that dictate the potential quality of the assembly. If WGS is initially being used for strain level identification with the option to be able to continue sequencing in order to generate an assembly of the genome then compromises may be required. In some cases it is not always practical to extract sufficient quantities of DNA to achieve this using traditional approaches.

Optimising a bead-based DNA extraction protocol for both improved yield and speed, we explored the effect of minimising the input and starting molecule length on nanopore performance to determine the speed and sensitivity of sequencing. We applied our optimised protocol to extract DNA, sequence and assemble the genome of a single *Drosophila* fly and show that a contiguous, high gene completeness assembly is possible.

## 2. Results

### 2.1. Comparison between spin columns and magnetic beads on DNA extraction metrics

We used equal amounts of the ZymoBIOMICS Microbial Community Standard to compare the effectiveness of precipitation and purification of DNA following bead beating in the presence of the Qiagen CD1 lysis solution, using both the Qiagen PowerSoil Pro spin columns and Kapa Biosystems magnetic beads. We compared both DNA yield and fragment length using the Life Technologies Qubit and an Agilent Genomic TapeStation. Yield for the PowerSoil Pro kit was > 355 ng of DNA in 50 μl (7.1 ng/μl) and for the Kapa beads was > 440 ng of DNA in 10 μl (44.1 ng/μl) with DNA Integrity Number (DIN) scores of 6.3 and 6.6, and average sizes of 14.4 and 14.0 Kbp respectively. Agilent Genomic TapeStation electropherograms for both DNA samples are shown in Figure 1.

**Figure 1:**
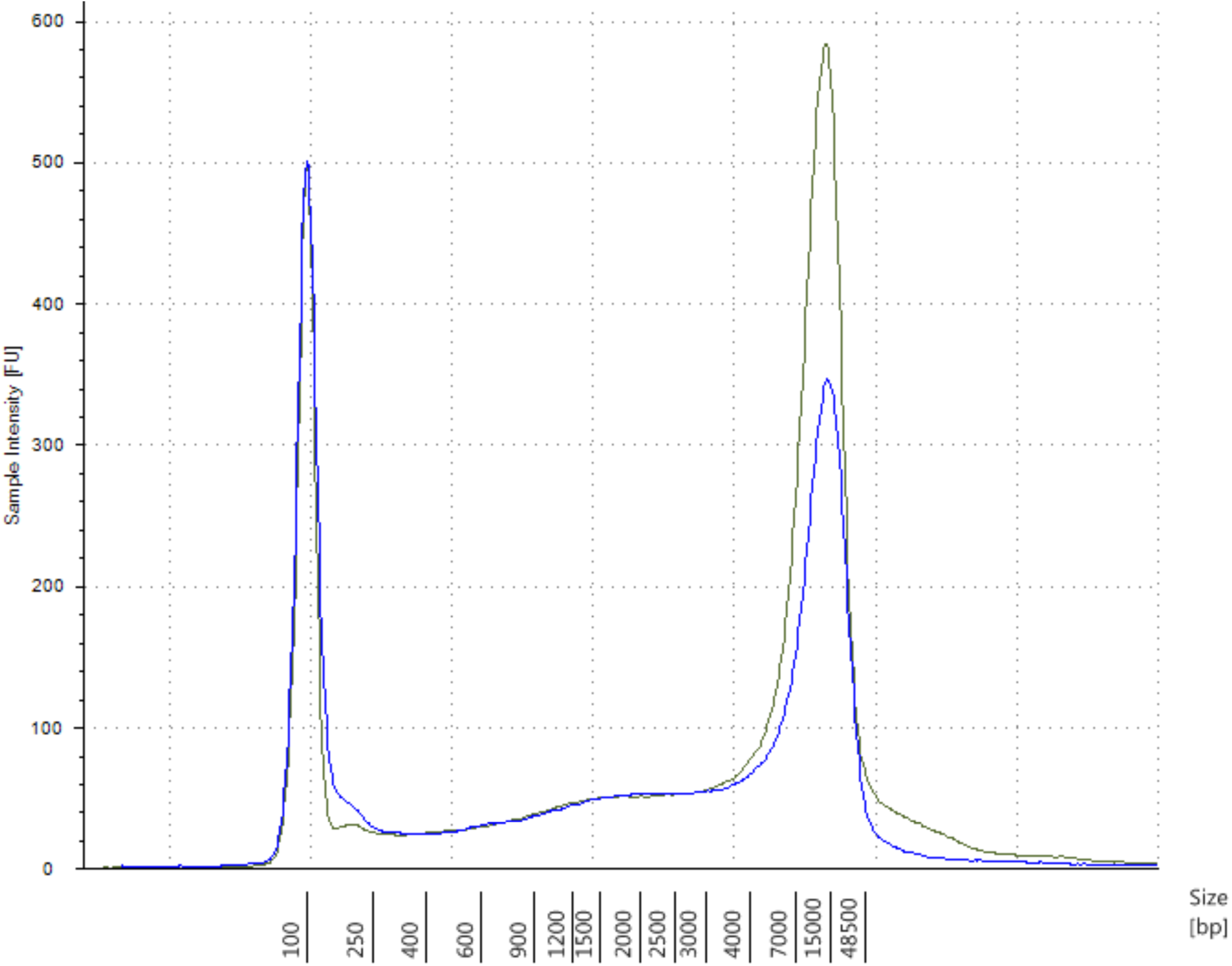
The difference between column and magnetic based DNA precipitation and purification methods. Overlaid Agilent genomic tapestation electropherograms of Poweroil Pro purified DNA (blue) and Kapa bead purified DNA (green).

### 2.2. Optimising DNA yield and molecule length

Having established that beads outperformed spin columns for both yield and molecule length we used this method to investigate the effect that the volume of lysis solution and bead beating weight had on these. Figure 2 shows the molecule lengths achieved with a range of bead weights and lysis volumes and Table 2 shows the corresponding DNA yields, average DNA molecule length according to the TapeStation and the resultant DIN.

**Figure 2:**
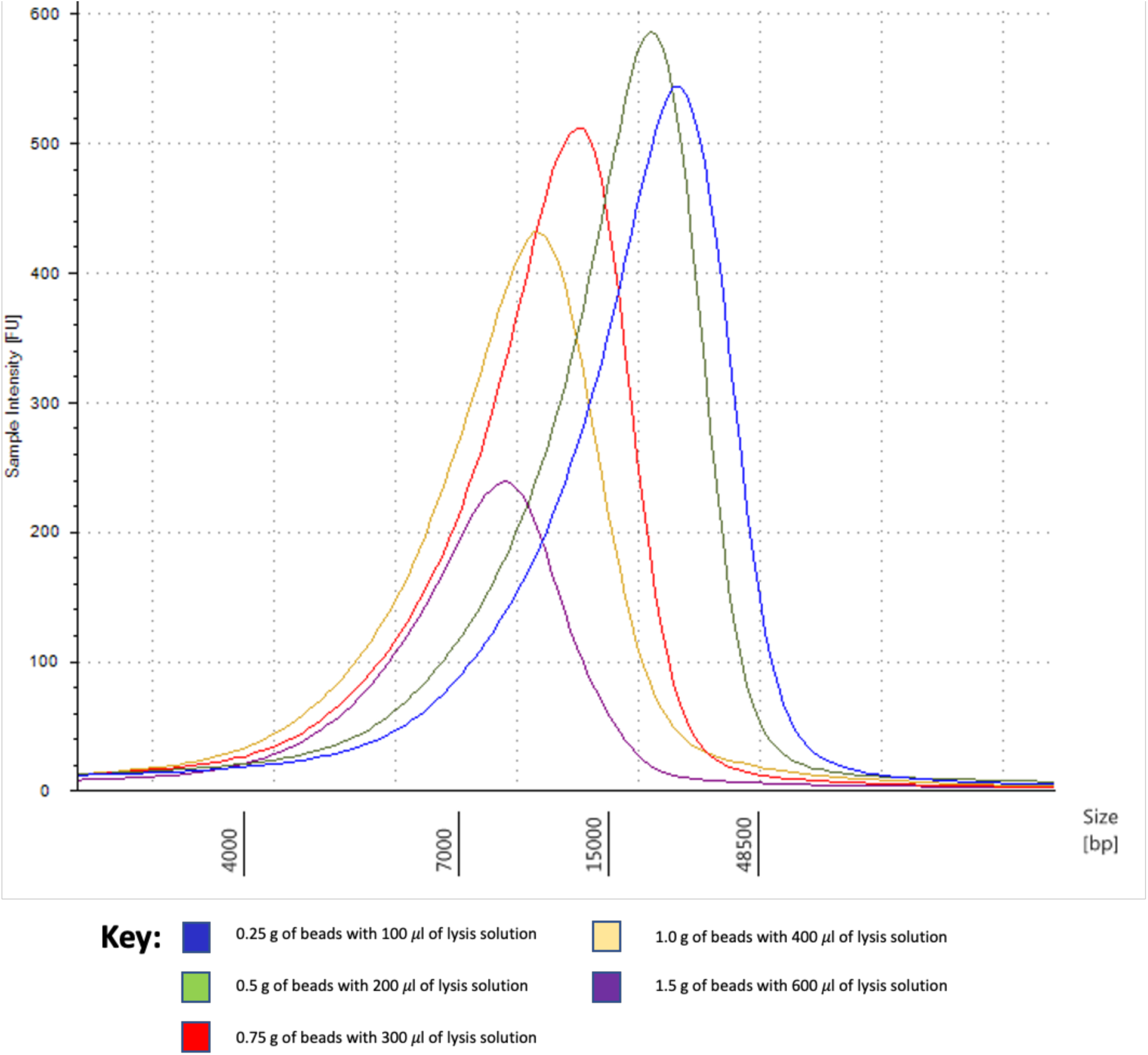
The effect of beating bead weight and lysis solution volume on DNA molecule length. Overlaid Agilent genomic TapeStation electropherograms of extractions from different combinations of lysis solution and bead weight.

**Table 1:** DNA extraction metrics. DNA yields, average molecule size and DNA integrity number (DIN) when using different beating bead weights and lysis solution volumes.

| Weight of beads (g) | Volume of lysis buffer (μl) | DNA yield (ng) | Average size (kbp) | Agilent DIN Scores |
| --- | --- | --- | --- | --- |
| 0.25 | 100 | 158 | 21.3 | 7.9 |
| 0.5 | 200 | 167 | 18.8 | 7.8 |
| 0.75 | 300 | 156 | 12.9 | 7.4 |
| 1 | 400 | 131 | 10.7 | 7.2 |
| 1.5 | 600 | 123 | 8.9 | 6.9 |

**Table 2:** Sequencing run metrics. Read N50 and read counts over time when reducing DNA input into Nanopore library construction using the ONT RAD004 kit.

| Experiment | Read N50 (bp) | Read count after 1 hr | Read count after 4 hrs | Read count after 8 hrs |
| --- | --- | --- | --- | --- |
| 200 ng input - full reaction | 4,014 | 140,753 | 671,323 | 1,365,480 |
| 100 ng input - full reaction | 4,079 | 95,725 | 471,984 | 930,413 |
| 50 ng input - full reaction | 3,899 | 60,772 | 295,950 | 562,340 |
| 10 ng input - full reaction | 4,032 | 9,005 | 34,241 | 55,651 |
| 1 ng input - full reaction | 3,907 | 834 | 3,327 | 6,119 |
| 25 ng input - half reaction | 2,788 | 73,998 | 388,397 | 756,514 |
| 10 ng input - quarter reaction | 2,683 | 33,786 | 164,660 | 312,868 |

### 2.3. Effect of DNA input and reaction volume on read N50 and read count

Having improved the yield and DNA molecule length we then turned our attention to reducing the input and reaction volumes. Using DNA extracted with the Kapa bead based protocol, described previously, we determined the number of reads generated over time for different DNA inputs and reaction volumes and calculated the read N50 (Table 2).

### 2.4. Single *Drosophila* DNA extraction and sequencing

Using the optimised magnetic bead and reduced bead beating weight and lysis volume protocol we wanted to determine whether this method would be suitable for in field whole genome sequencing of low DNA yielding species.

To replicate this type of study we used a single *Drosophila* individual and achieved a DNA yield of > 150 ng DNA with a molecule length around 6 Kbp according to the Agilent TapeStation (Figure 3).

**Figure 3:**
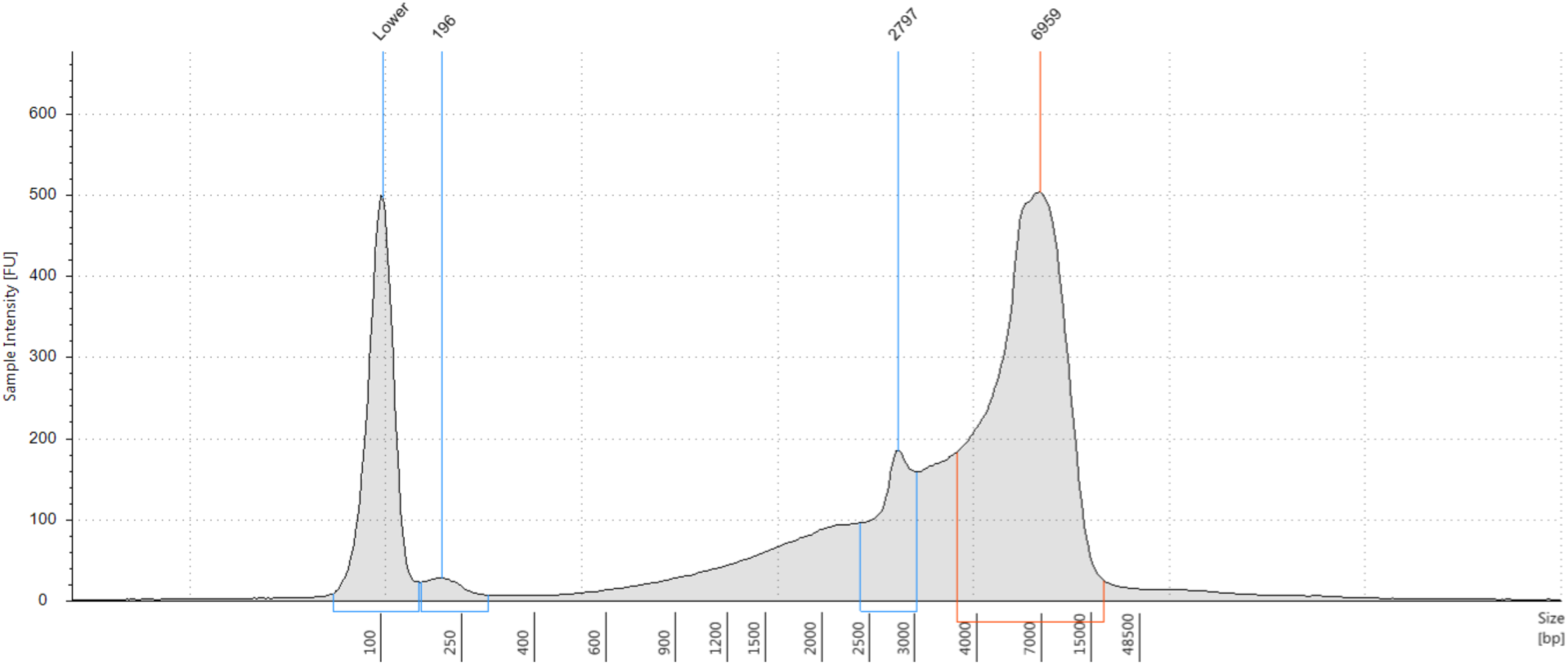
Molecular weight determination of the single Drosophila DNA extraction. Agilent Genomic Tape Electropherogram showing the range of DNA molecule lengths from the bead beaten single Drosophila DNA extraction.

After DNA QC library construction was performed with 110 ng of DNA in half the recommended ONT RAD004 reaction volume. Nanopore sequencing resulted in > 3.75 million reads with an N50 of 3.12 Kbp and a total yield of > 7.5 Gbp. Read length distribution for this run is shown in Figure 4.

**Figure 4:**
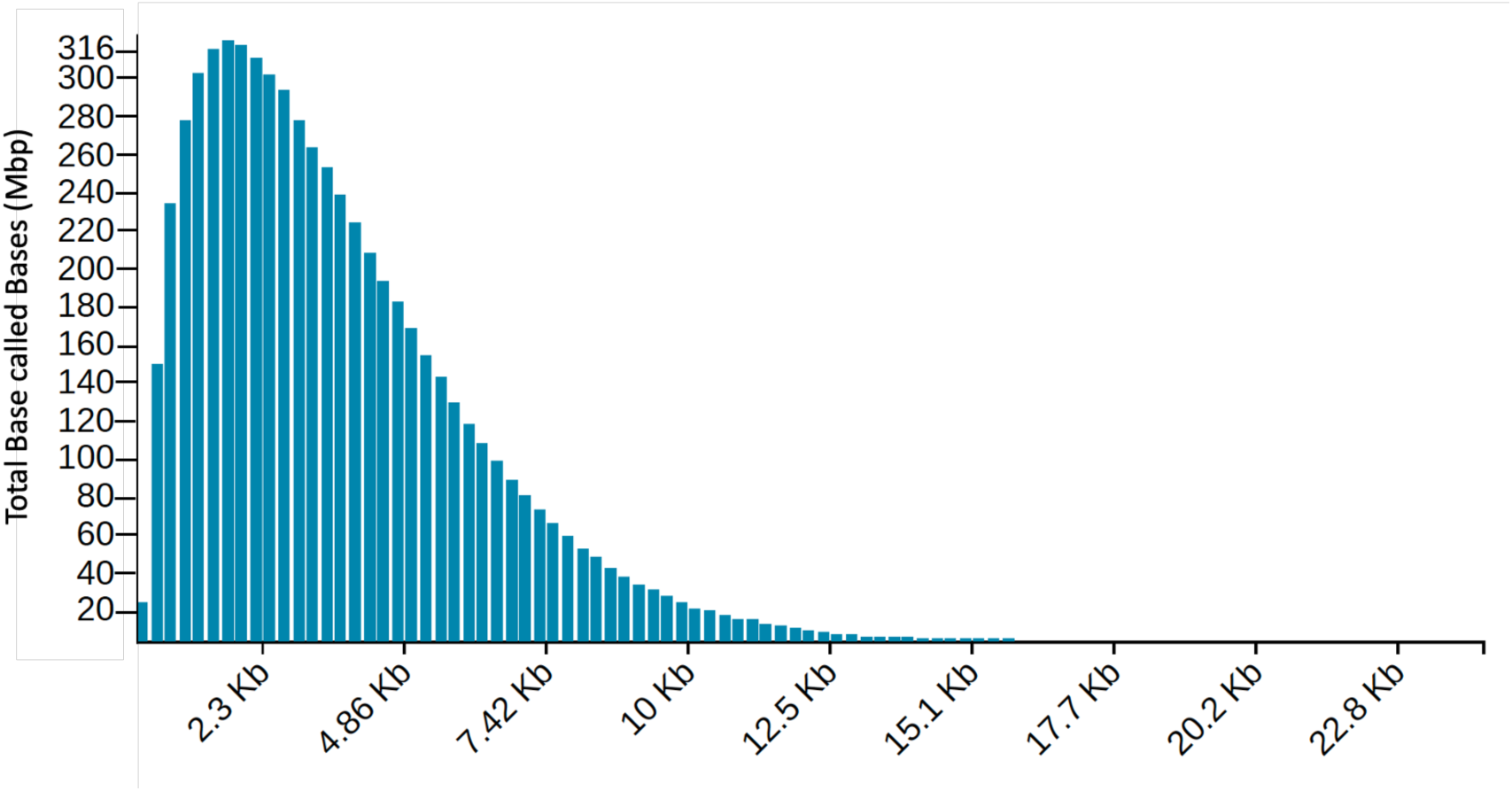
Nanopore read length distribution. Plot from the MinKNOW run report showing the distribution of read lengths generated when sequencing the single Drosophila.

### 2.5. Single *Drosophila* assembly and analysis

After 36 hours of sequencing, pore performance had dropped and there was no appreciable increase in sequence outputs. We performed assembly on the data generated at cumulative 6 hour intervals up to 36 hours and calculated the number of contigs, total length of the assembly, contig N50 and contig count (Table 3). By 36 hours of sequencing, half of the genome was composed of 31 contigs of at least 1.23 Mb. However, even after 12 hours of data generation, half the genome was contained in only 48 contigs.

**Table 3:** Single Drosophila genome assembly metrics at various time points of sequence collection. Assembled contigs, assembled length, contig N50 and contig N50 count after 6, 12, 18, 24, 36 and at the end of sequencing.

| Data collection time (h) | Contigs | Total Length (bp) | N50 (bp) | N50 Count |
| --- | --- | --- | --- | --- |
| 0 - 6 | 2,875 | 111,298,559 | 96,988 | 328 |
| 0 - 12 | 1,926 | 123,564,754 | 624,307 | 48 |
| 0 - 18 | 1,751 | 125,181,682 | 890,178 | 39 |
| 0 - 24 | 1,720 | 124,760,854 | 968,324 | 37 |
| 0 - 30 | 1,738 | 124,725,349 | 1,102,489 | 33 |
| 0 - 36 | 1,755 | 125,006,991 | 1,237,631 | 31 |
| all reads | 1,622 | 125,192,471 | 1,237,646 | 31 |

We used QUAST to compare our assembly to the gold standard reference sequence and also to the hybrid Illumina/Nanopore/Hi-C assembly described previously. This showed our assemblies after 18 hours represented > 98% of the genome fraction of the hybrid assembly, but around 85% of the gold standard (Table 4). To check the gene completeness of our assemblies, we performed BUSCO analysis on each (Table 4). Assembled contigs were then aligned to the Gold Standard reference genome using ALVIS (Figure 5).

**Table 4:** Single gene orthologue metrics and assembly comparison. Analysis of the assembled genome showing the percentage of complete, fragmented and missing BUSCO genes and QUAST comparison against the Gold Standard Reference and a Hybrid assembled genome.

| Seq'ing Time | BUSCO |  |  | Gold Standard Reference |  |  | Hybrid Assembly |  |  |
| --- | --- | --- | --- | --- | --- | --- | --- | --- | --- |
|  | Complete | Fragmented | Missing | Genome Fraction | Misass-embles | NGA50 | Genome Fraction | Misass-embles | NGA50 |
| 0h-6h | 85.7% | 2.1% | 12.2% | 76.0% | 1744 | 40,431 | 88.6% | 2915 | 57,476 |
| 0h-12h | 96.1% | 0.7% | 3.2% | 84.4% | 2105 | 99,655 | 97.5% | 3794 | 153,384 |
| 0h-18h | 96.7% | 0.5% | 2.8% | 85.5% | 2269 | 118,821 | 98.2% | 4170 | 163,047 |
| 0h-24h | 96.6% | 0.6% | 2.8% | 85.2% | 2218 | 120,572 | 98.3% | 4139 | 163,798 |
| 0h-30h | 96.6% | 0.6% | 2.8% | 85.2% | 2221 | 120,416 | 98.2% | 4034 | 166,127 |
| 0h-36h | 96.8% | 0.6% | 2.6% | 85.3% | 2274 | 120,275 | 98.3% | 4082 | 163,785 |
| all reads | 96.9% | 0.5% | 2.6% | 85.5% | 2308 | 122,859 | 98.3% | 4136 | 166,559 |

**Figure 5:**
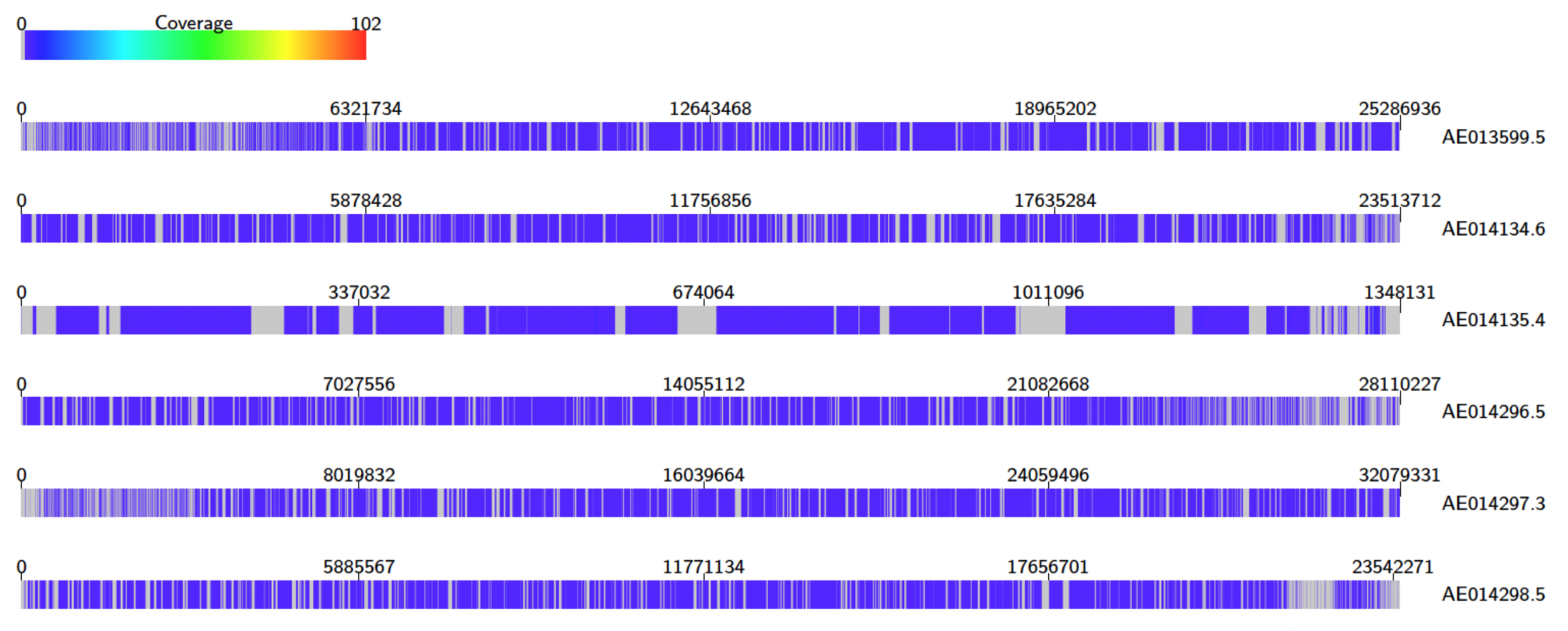
ALVIS alignment of the single Drosophila assembled contigs versus the Gold Standard Reference. Alignment of the assembled contigs against Chromosomes 2R (AE013599.5), 2L (AE014134.6), 4 (AE014135.4), 3L (AE014296.5), 3R (AEAE014297.3) and X (AEAE014298) of the reference genome.

## 3. Methods

### 3.1. DNA extractions

DNA was extracted from the ZymoBIOMICS Microbial Community Standard cells (Zymo Research, Irvine, CA, USA) using the CD1 lysis buffer and beating beads from a Qiagen PowerSoil Pro Kit (Qiagen, Hilden, Germany) and then either the remaining components of the PowerSoil Pro kit or DNA purification beads from Kapa Biosciences (Roche, Burgess Hill, UK) and either the MPBio SuperFastPrep-2 (MPbio, Eschwege, Germany) or SPEX Sample Prep 2010 GenoGrinder (SPEX Europe, Rickmansworth, UK).

### 3.2. Comparison between spin columns and magnetic beads on DNA extraction metrics

Two 2 ml tubes containing 1.5 gs of PowerSoil Pro beads and 600 *μ*l of CD1 buffer and 10 *μ*l of ZymoBIOMICS Microbial Community Standard cells were placed in the SuperFastPrep-2 and mixed for 20 seconds at a speed code of 20. The tube was then spun for 30 seconds at 10,000 rcf in an Eppendorf 5415R centrifuge (Eppendorf, Stevenage, UK) and the supernatants pooled in a fresh 1.5 ml Lobind Eppendorf tube (Eppendorf).

For the PowerSoil Pro kit test, 200 *μ*l of Solution CD2 was added to 550 *μ*l of lysed and bead beaten cells and vortexed for 5 seconds before spinning at 10,000 rcf for 1 minute. The supernatant was then transferred to a clean 2 ml Microcentrifuge Tube along with 600 *μ*l of Solution CD3 and vortexed for 5 seconds. A 650 *μ*l aliquot of this lysate was then applied to theMB Spin Column, centrifuged at 10,000 rcf for 1 minute and the flow through discarded. This step was repeated to ensure that all of the lysate has passed through the MB Spin Column. To wash the column 500 *μ*l of Solution EA was added to the MB Spin Column and centrifuged at 10,000 rcf x g for 1 minute with the flow through discarded and this was repeated with 500 *μ*l of Solution C5. The column was placed into a clean tube, spun for 2 minutes at 10,000 rcf and then placed in a fresh 1.5 ml Lobind tube and 50 *μ*l of Solution C6 was added to the center of the white filter membrane. This was left at room temperature for 5 minutes before being centrifuged at 10,000 rcf for 1 minute to elute and collect the DNA.

For the magnetic bead test 550 *μ*l of Kapa beads was added to 550 *μ*l of lysed and bead beaten cells, vortexed and then incubated at room temperature for 3 minutes. The tube was then pulse spun in a microfuge then placed in a magnetic particle concentrator and the beads allowed to concentrate. The supernatant was discarded and the beads were washed twice with fresh 70% ethanol. Care was taken to remove all the ethanol and the tube removed from the MPC and the beads resuspended in 10 μl of Qiagen CD6 buffer and incubated at room temperature for 2 minutes. The tube was then pulse spun in a microfuge then placed in a magnetic particle concentrator and the beads allowed to concentrate. The supernatant containing the DNA was then transferred to a fresh 1.5 ml Lobind Eppendorf tube.

For both methods a 1 *μ*l aliquot of DNA was used to determine concentration using the Qubit HS assay (Life Technologies, Loughborough, UK) and a second 1 *μ*l aliquot was used to determine molecule length with an Agilent Genomic Tape (Agilent, Cheadle, UK) on an Agilent TapeStation (Agilent).

### 3.3. Optimising DNA yield and molecule length

In a 2 ml tubes multiples of 0.25 g of powersoil pro beads and 100 *μ*l of CD1 buffer and 2 *μ*l of ZymoBIOMICS Microbial Community Standard cells added (see Table 1) then placed in the GenoGrinder and mixed for 5 minutes at 1,500 rpm in a 2 ml well block. The tube was then spun for 30 seconds at maximum speed in an Eppendorf 5427R centrifuge and the supernatant transferred to a fresh 1.5 ml Lobind Eppendorf tube. An equal volume of Kapa beads was added, vortexed and then incubated at room temperature for 3 minutes. The tube was then pulse spun in a microfuge then placed in a magnetic particle concentrator and the beads allowed to concentrate. The supernatant was discarded and the beads were washed twice with fresh 70% ethanol. Care was taken to remove all the ethanol and the tube removed from the MPC and the beads resuspended in 10 *μ*l DNase free water and incubated at room temperature for 2 minutes. The tube was then pulse spun in a microfuge then placed in a magnetic particle concentrator and the beads allowed to concentrate. The supernatant containing the DNA was then transferred to a fresh 1.5 ml Lobind Eppendorf tube. A 1 *μ*l aliquot was used to determine DNA concentration using the Qubit HS assay and a second 1 *μ*l aliquot was used to determine DNA molecule length with an Agilent Genomic Tape on an Agilent TapeStation.

### 3.4. *Drosophila melanogaster* DNA extraction

DNA was extracted from a single *Drosophila melanogaster* fly using the lysis buffer and beating beads from a Qiagen PowerSoil Pro Kit, DNA purification beads from Kapa Biosciences and a MP Biomedical SuperFastPrep-2.

In a 2 ml tube 0.25 g of PowerSoil Pro beads, 100 *μ*l of CD1 buffer and a single fly were placed in the SuperFastPrep-2 and mixed for 20 seconds on a speed setting of 25. DNA was precipitated and washed with ethanol as described previously except that DNA was eluted from the Kapa beads in 5 *μ*l of DNase free water rather than 10 *μ*l and QC was performed on 0.5 *μ*l aliquots rather than 1 *μ*l.

### 3.5. Effect of DNA input and reaction volume on read N50 and read count

We investigated the effect of reducing DNA input amounts and reaction volumes using DNA extracted from the ZymoBIOMICS Microbial Community Standard cells and the ONT SQK-RAD004 kit. DNA input amount, reaction volumes and amount of RAP adapter added for each experiment is shown in Table 5.

**Table 5:** Variable DNA input reaction metrics. DNA input, reaction volume and the amount of RAP adapter added in testing the effect of input on sequencing metrics.

| Experiment | Transposase reaction volume (μl) | RAP adapter added (μl) |
| --- | --- | --- |
| 200 ng input | 10 | 1 |
| 100 ng input | 10 | 1 |
| 50 ng input | 10 | 1 |
| 10 ng input | 10 | 1 |
| 1 ng input | 10 | 1 |
| 25 ng input- half reaction volume | 5 | 0.5 |
| 10 ng input- quarter reaction volume | 2.5 | 0.25 |

The initial library mix was incubated for 1 minute at 30 °C followed by 1 minute at 80 °C, the reaction cooled to room temperature and the appropriate amount of the RAP adapter added. This was vortexed to mix and the reaction incubated at room temperature for 5 minutes.

If required, the transposase plus RAP adapter reaction volume was then made up to 11 *μ*l with EB, 34 *μ*l of SQB, 25.5 *μ*l of LBB and 4.5 *μ*l of DNase free water added, then mixed and loaded onto a primed 9.4.1 ONT MinION flowcell according to the manufacturer’s instructions. Reads after 1, 4 and 8 hours were calculated and read length N50 recorded.

### 3.6. Single *Drosophila* sequencing

For the single *Drosophila* extraction the SQK-RAD004 kit was used and 110 ng of DNA in 3.75 *μ*l volume was mixed with 1.25 *μ*l of the FRA and incubated for 1 minute at 30°C followed by 1 minute at 80 °C, the reaction was then cooled to room temperature and 0.5 *μ*l of the RAP adapter was added, vortexed and the reaction incubated at room temperature for 5 minutes. To this 34 *μ*l of SQB, 25.5 *μ*l of LB, 5.5 *μ*l of EB and 4.5 *μ*l of DNase free water were added then mixed and loaded onto a primed 9.4.1 ONT MinION flowcell according to the manufacturer’s instructions.

### 3.7. Single *Drosophila* assembly

Reads were basecalled with Guppy v5.0.7 using the “super accuracy” basecalling model (dna_r9.4.1_450bps_sup). All reads that passed Guppy’s own QC were collected and placed into bins depending on the time of sequencing using a custom script (available in the repository https://github.com/SR-Martin/How-low-scripts). Read-sets for each window 0-n hours (n=1,2,3,4,5,6) were created by manually combining bins. Each read-set was assembled using Flye v2.8.1^[25]^ with the options --nano-raw and --genome-size 180m.

Each assembly was compared to the “Gold Standard Reference”^[26]^ and the “Hybrid Assembly” using QUAST v5.0.2 with default parameters. Each assembly was also assessed using BUSCO v4.1.2 against the database diptera_odb10 and the assembled contigs aligned to the “Gold Standard Reference” using ALVIS^[27]^.

## 4. Discussion

In our experiments we were able to show that our straightforward magnetic bead based DNA recovery rates were > 20% greater than the traditional column based methods and we were able to optimise the protocol further to improve DNA molecule length and quality. We demonstrated the effect of reducing input on ONT sequencing performance and then applied these findings to generate a genome assembly from a single *Drosophila* identifying > 96% of BUSCO genes as complete. The methods that we optimised were designed to be transportable and allow for remote DNA extraction and sequencing.

For the bead weight and lysis volume test we switched to the Spex GenoGrinder from the SuperFastPrep-2 to ensure that each of the tubes experienced exactly the same bead beating time and shaking. It is capable of processing up to 96 samples at a time in 2 × 48 well blocks rather than the SuperFastPrep-2 which can only handle 2 tubes at a time. Interestingly our results suggested that reducing the bead weight and lysis volumes actually increased DNA quantity and quality. We observed average sizes increase from 8.9 to 21.5 Kbp and DIN scores from 6.9 to 7.9. In our experience the TapeStation can overestimate molecule length but we would expect that the DNA extracted using 100 *μ*l of lysis solution combined with 0.25 g of beads would generate reads with an average size > 10 Kbp and an N50 in the region of 15 Kbp if sequenced on a flowcell using the 1D ligation chemistry. It was encouraging to note that the DIN number for the extraction of the 600 *μ*l lysis solution plus 1.5 g of beads was close to that observed in the column versus Kapa beads test (6.9 to 6.6) and the reduction in yield (123 ng v 440 ng) not dissimilar taking into account the reduction in ZymoBIOMICS Microbial Community Standard cells used (2 v 10 *μ*l). This highlights the robustness of the protocol and scalability with respect to the starting concentration of material.

For testing our DNA extraction protocols we chose the ZymoBIOMICS Microbial Community Standard as it contains 5 Gram-positive bacteria, 3 Gram-negative bacteria and 2 yeasts with genome sizes ranging from 1.9 to 18.9 Mbp and GC contents from 32.9 to 66.2% and is therefore a good test for method development and comparisons. Whilst bead beating was expected to help with DNA extractions from yeast, which can be notoriously difficult to handle, improved DNA yields seen when reducing bead weight and lysis volumes indicates that there is no loss of sensitivity in doing so. In fact the data suggests that this might result in a more representative extraction as DNA yields were greater.

When investigating the effect of reducing DNA input we observed 5x the number of reads after 4 hours and 10x the number of reads after 8 hours compared to 1 hour for the 50 to 200 ng input full reaction, the 25 ng input half reaction and the 10 ng input quarter reaction. This suggests that neither the rapid DNA extraction protocol or the reduced DNA input negatively affected flowcell performance. Flowcell performance did drop off when using < 50 ng in a full reaction volume reaction indicating that this combination could be a limiting step if only loading a single sample.

It has been reported that when using transposases for Illumina sequencing the ratio of DNA to transposase can be adjusted to alter fragmentation sizes. In our experiments when reducing the input amounts and maintaining reaction volumes we didn’t observe any increase in read length N50. With the full reaction volumes our read N50 remains consistent ranging between 3.9 and 4.1 Kbp. This suggests that the amount of transposase in these kits might be limiting. Calculating that 400 ng of DNA at 10 Kbp equates to > 3.9 × 10^10^ individual strands of DNA and 10 ng has > 9.7 × 10^8^, at most you might expect to achieve 5 million sequence reads on a flowcell there is a vast excess of molecules present. Whilst this excess may be necessary to ensure efficient sequencing it might not be necessary to have them all as viable library molecules capable of being sequenced.

When we reduce the reaction volume we observe a reduction in read N50 length. With 25 ng in a half reaction volume and 10 ng in a quarter we observe read N50s of 2.8 and 2.6 Kbp respectively. Given that in these two cases the reaction volume is halved and the inputs roughly halved this supports the hypothesis that the transposase might be a limiting factor. For rapid diagnostic applications shorter reads are beneficial as they pass through the pores more quickly freeing them up for the next molecule to enter. This is borne out when we look at the number of reads generated for the 50 ng input full reaction volume versus the 25 ng half reaction volume. For the 50 ng experiment 60 K reads are generated in the first hour compared to 75 K for the 25 ng input. This equates to a 25% increase and can be explained in part by the shorter read N50 (2.7 v 3.9 Kbp) resulting in more reads being processed during this time. As ONT improves their base calling accuracy there will be more scope to reduce read lengths and we expect to see further advancement in this area.

When applying our optimised DNA extraction protocol to a single *Drosophila* we achieved a read N50 of 3.12 Kbp. By modern extraction methods this figure may be deemed very low. However, close analysis of the read distribution plot shows that there are >11,000 reads > 10 Kbp which total > 125 Mbp of sequence and overall we achieved > 50x genome coverage. Assembling the data with Flye resulted in a BUSCO completeness score of 96.9% (it is 98% in the Gold Standard Reference) and with a total assembled content in excess of 125 Mbp representing 85.5% of the reference genome. It could be argued that we might have achieved a better assembly with a ligation based approach but with 110 ng of input it is unlikely that any sequencing run would have been as successful in generating sufficient genome coverage to take advantage of the longer read lengths. Ligation methods also require more time, more equipment and a robust cold chain to be implemented.

Our assembly compared favourably against a much more expensive triple hybrid assembly which employed Illumina, Nanopore and Hi-C data, with our assembly representing ∼ 98% of its content. The ALVIS alignment against the Gold Standard Reference genome was also encouraging. Each of the assembled contigs only aligned to a single place on the genome suggesting that the missing components from the genome are repeat structures greater than those that we could resolve given the read lengths that we achieved. It is worth noting that the total consumable cost of generating our sequence data including DNA extraction was < £600.

Bead beating is widely used for metagenomic analysis as it maximises the number of different species that have their cell walls ruptured releasing DNA. Improving the molecular weight of extracted DNA will have a positive effect on the ability to obtain complete MAGs. Our finding that DNA molecule length can be improved by adjusting and reducing the weight of beads used and lysis volume could be beneficial in a range of projects and has the potential to be further refined. When used in conjunction with 1D ligation chemistry this should result in improved sequence reads lengths which in turn should result in improved assembly metrics. Longer DNA molecules also improve the efficiency of adaptive sampling on nanopore flowcells^[28]^. The longer the DNA molecules, the better the enrichment of less abundant species and this in turn will lead to better species classification and MAGs.

In time, the need to bead beat might be countered by the discovery of further enzymes capable of digesting cell walls, such as those used in MetaPolyzyme, and this is an active area of research but for many, beads will still provide the best option for metagenomic DNA extraction due to the speed of processing samples and yield benefits. For scientists working with low biomass samples such as air and clinical samples such as fine needle aspirations or core needle biopsies, improved DNA recovery rates combined with the ability to reduce the amount of DNA required for successful sequencing will be beneficial. To overcome low yields many employ amplification techniques such as whole genome amplification (WGA) to bulk up the amount of material they have. However, the network of branches formed during WGA is known to block pores and reduce pore life in nanopore flowcells so it is recommended that these sample types be debranched using enzymes such as T7 or S1 that target and nick single stranded stretches. With modern WGA techniques taking a minimum of 90 minutes and debranching adding further time to process a sample ahead of sequencing this would delay the time to diagnosis for clinical samples.

One of the aims of our study was to reduce both the amount of input material required and the time to generate results. In previous studies clinically significant data has been identified with as little as 13,000 sequence reads^[3]^. Our optimised DNA extraction protocol is capable of yielding sufficient material for sequencing within 10 minutes of sample collection. Combining this with the 10 minute rapid library prep requiring only 25 ng of input material, we demonstrate that enough data can be generated within 45 minutes of sample collection to reach the necessary sequence read number. Scientists looking to assemble genomes, such as those associated with the EBP, and are challenged by low DNA yielding species, will be encouraged by these results. Our protocols were developed with field work in mind. Using the handheld SuperFastPrep-2 showcased its ability in our DNA extraction pipeline and with devices such as ONT’s Mk1C and MiniPCR available alongside improved temperature stability of MinION flowcells and lyophilised field sequencing kits, all of our protocols can easily be translated into methods that can be applied by anyone, anywhere.

These robust methods are capable of maximising DNA yield and in requiring less DNA it is easy to envisage a scenario whereby a sample can be collected in the field, have its DNA extracted and sequenced there and then. The data can be analysed in real time and at the point of collection and once it can be determined if further sequencing is required and a decision made to continue with sequencing or to perform a nuclease flush to ready the flowcell for the next sample. It might be that assemblies like we achieved for *Drosophila* may not be sufficient for the project in which case DNA could be extracted from part of the body preserving some to be taken back to the laboratory for a higher quality DNA extraction or if the sample is so small can dictate whether additional specimens are required.

## 5. Conclusions

In optimising the DNA yield and molecule length achieved through bead beating we have developed a simple and rapid yet robust method for DNA extraction. Combining this with redefining the DNA input requirements for transposase based nanopore sequencing we have built a low input, versatile pipeline capable of sequencing samples from a wide variety of sources and settings. Sequencing can be initiated within 20 minutes of sample collections and we have demonstrated its ability to produce high quality gene complete assemblies for consumable sequencing costs of less than £600.

## Ethical Approval

Not applicable

## Consent for publication

Not applicable

## Availability of data and metadata

The sequence datasets generated and analysed during the current study are available in the European Nucleotide Archive (http://www.ebi.ac.uk/ena) repository under accession number PRJEB48132.

## Author contribution

DH, RML, RM and MDC designed the study. DH, DC, MG and PC performed the experiments. MA, SM, and RML performed data analysis. DH, RML, DC, PC, PA, MG, MA, SM, RM and MDC wrote and edited the manuscript.

## Acknowledgements

We would like to thank Will Nash at the Earlham Institute for providing the *Drosophila* sample used in this study.

This research was supported in part by the NBI Computing Infrastructure for Science Group, which provides technical support and maintenance to Earlham Institute’s high-performance computing cluster and storage systems.

